# 3D cell culture stimulates the secretion of in vivo like extracellular vesicles

**DOI:** 10.1101/556621

**Authors:** Sirisha Thippabhotla, Cuncong Zhong, Mei He

## Abstract

For studying cellular communications ex-vivo, a two-dimensional (2D) cell culture model is currently used as the “gold standard”. 2D culture models are also widely used in the study of RNA expression profiles from tumor cells secreted extracellular vesicles (EVs) for tumor biomarker discovery. Although the 2D culture system is simple and easily accessible, the culture environment is unable to represent in vivo extracellular matrix (ECM) microenvironment. Our study observed that 2D culture-derived EVs showed significantly different profiles in terms of secretion dynamics and essential signaling molecular contents (RNAs and DNAs), when compared to the three-dimensional (3D) culture-derived EVs. By performing small RNA next-generation sequencing (NGS) analysis of cervical cancer cells and their EVs compared with cervical cancer patient plasma EV-derived small RNAs, we observed that 3D culture-derived EV small RNAs differ from their parent cell small RNA profile which may indicate a specific sorting process. Most importantly, the 3D culture derived EV small RNA profile exhibited a much higher similarity (~96%) to in vivo circulating EVs derived from cervical cancer patient plasma. However, 2D culture derived EV small RNA profile correlated better with only their parent cells cultured in 2D. On the other hand, DNA sequencing analysis suggests that culture and growth conditions do not affect the genomic information carried by EV secretion. This work also suggests that tackling EV molecular alterations secreted into interstitial fluids can provide an alternative, non-invasive approach for investigating 3D tissue behaviors at the molecular precision. This work could serve as a foundation for building precise models employed in mimicking in vivo tissue system with EVs as the molecular indicators or transporters. Such models could be used for investigating tumor biomarkers, drug screening, and understanding tumor progression and metastasis.

## Introduction

Extracellular vesicles (EVs) are a heterogeneous mixture of membranous structures, derived from either the cellular endosomal pathway or shed from the plasma membrane^1^. Due to the different biogenesis, EVs are generally classified as the exosomes (~30-250 nm) and microvesicles (~100-1000nm)^2–5^. Microvesicles with a particle size up to 1 μm are formed by the outward budding from the cell membrane ^1, 6^. In contrast, exosomes are one subgroup of EVs formed from the intercellular structure via the inward invagination of endosome membranes^7^, which encapsulated complex cellular signaling components, including proteins, lipids, and nucleic acids (e.g., mRNA, miRNA, DNA)^8, 9^. However, the current definition of exosomes is still not well defined. Assigning an EV to a particular biogenesis pathway remains extraordinarily difficult. In order to be consistent with the International Society for Extracellular Vesicles (ISEV community), we use the generic term “extracellular vesicles (EVs)” while referring to small vesicles (< ~250 nm)^10^. EVs play important roles in cell-to-cell communications via cross-transferring of important signaling components^11–14^. For instance, EVs can carry and transmit mRNAs and miRNAs for promoting tumor growth, which has been identified as the diagnostic biomarkers for lung cancer^15^, breast cancer^16^ and ovarian cancer^17, 18^. The mRNAs found in smaller EVs (exosomes) can be transferred and translated to functional proteins after uptake by other cells ^19^. The RNA content in EVs varies depending on the cell type and cellular status from their parent cells, for performing cell-to-cell communications^20–25^. The mechanism of how specific RNA sequences are selectively packaged into EVs is still unknown, but is an intensive area of investigation^3, 26–28^.

For investigating cellular communications and behaviors ex vivo, presently, the two-dimensional (2D) cell culture model is widely used as the “gold standard”^29^. This 2D culture system serves as an essential model for investigating tissue physiology and complex biological activity, from cell differentiation to tissue morphogenesis^30^. Many 2D cell culture systems have also been widely employed for studying EV RNA expression profiles from tumor cells ^31^, roles in promoting tumor growth ^17^, and tumor biomarker discovery ^32–34^. Although the 2D culture system provides simple cell attachment and nutrients supply, the flat and hard surface from plastic or glass substrates are unable to represent the in vivo extracellular matrix (ECM) microenvironment in tissue or organs^35^. The monolayer cells under 2D culture condition completely differ from in vivo status where cells grow in three dimensions (3D), in terms of cell morphology, cell-to-cell interactions, growth behavior, and interactions with extracellular matrix^30^. It has been well demonstrated that 2D cell monolayer is unable to represent the physiology of in vivo 3D tissues or organs, due to the substantially different microenvironment (e.g., mechanical and biochemical properties) in tissue architecture^36–41^.

Unlike the 2D culture, 3D cell culture is more recognized for mimicking in vivo cellular behavior^41^. Instead of cell-to-cell interaction only by the edge in the 2D culture system, 3D cell culture involves cellular stretch and interactions from all angles, as well as the cell-to-ECM interactions^35^. These interactions aid in promoting cellular signaling transduction and proliferation. For instance, the behaviors of 3D tumor spheroids are more clinically relevant, considering hypoxia in the drug screening research^42^. Skin cells under 3D culture conditions survive better than 2D conditions when exposed to cytotoxic agents^43^. The ECM serves as the scaffold which is critical for cell adhesion, movement, stretch, proliferation and differentiation, and the prevention of apoptosis^44^. Within the 3D culture environment, we observed that cellular EV secretion dynamics and molecular contents are altered, compared to the 2D culture derived EVs. This observation indicates that 3D culture derived EV small RNAs could reflect in vivo tissue-derived EV RNAs. Although studies have been performed on EV secretion from human plasma using a 2D culture sytem^45, 46^, there have been no reports on the EV secretion landscape from 3D culture and their potential for mimicking in vivo tissue system. The EV biogenesis mechanisms, molecular sorting, vesicle trafficking and releasing are still not well understood so far^7, 26^. Other reports and our observation support that EV secretion behavior and molecular cargoes are altered by the influence of many factors including parent cell types, physiological and pathological status, and the stimuli from microenvironments^47^. Thus, only EVs derived from parent cells under biomimetic tissue culture conditions can reflect the in vivo EVs, which is extremely essential for studying cellular biological function, cell-to-cell communications and signaling transductions accurately. Such discoveries could eventually lead to the development of clinically significant EV biomarkers and therapeutic agents.

In this paper, we observed that 2D culture derived EVs showed significantly different profiles in terms of secretion dynamics and essential signaling molecular contents (RNAs), compared to the 3D culture derived EVs. By next-generation sequencing (NGS) analysis of cervical cancer cells and cervical cancer patient plasma-derived EV RNA, we observed that 3D culture derived EV small RNAs may undergo specific sorting process that differs from their parent cells. Most importantly, the 3D culture derived EV small RNA profile exhibited a much higher similarity (~ 96%) to in vivo circulating EVs from cervical cancer patient plasma, in comparison with 2D derived EV small RNA profile. On the other hand, DNA sequencing analysis suggests that culture and growth conditions do not affect the genomic information carried by EV secretion. By tackling EV molecular contents and alterations collected from interstitial fluids, we can provide an alternative, non-invasive approach for investigating 3D tissue behaviors at the molecular precision. This work could serve as a foundation for building precise model employed in mimicking in vivo tissue system, with EVs as the molecular indicators or vehicles, for investigating tumor biomarkers, drug screening, and understanding tumor progression and metastasis.

## Results

### EV secretion dynamics from 3D cultured cells significantly differ from 2D culture system

In order to culture in-vivo like 3D cells, we choose to use peptide hydrogel as the scaffolding material which has high biocompatibility and self-healing property for transforming between liquid and gelation^48–50^. This peptide hydrogel is stable in neutral pH and remains as gel formation in a wide range of temperatures from 2° C to 80° C^51, 52^. We examined the peptide hydrogel scaffolding structures in Figure 1 A to evaluate the 3D ECM microenvironment, which shows the dense mesh network with pore size nearly around 500 nm. The flexible mechanical tensile of peptide gel allows cells to proliferate and expand as a 3D spheroid. Due to the high biocompatibility, the big 3D spheroid still can attach well with the scaffolding support. We observed four classic morphologies presented in the 3D cultured spheroids with peptide hydrogel scaffolds: round, mass, grape, and stella, as shown in Figure 1B, which is consistent with the literature report^53, 54^. Confocal fluorescence image was further used to confirm the round tumor-like morphology in Figure 1C, which exhibits the typical in vivo tumor-like tissue architecture^55^ with each round cells laid over to form tumor spheroid. Figure 1C is a 3D confocal imaging reconstruction of cervical tumor cell spheroid using our 3D culture method. The heat map encoded red to blue color is to display the z-dimension change (height of tumor spheroid) which can give a better visualization of tumor cell distribution in the spheroid. The cervical cancer HeLa cells formed round spheroids after 4-5 days culture which are the most typical tumor morphology dominated in our 3D culture system. The grape and stellate shapes were observed occasionally. In strong contrast, 2D cultured HeLa cells are in the flat spreading and attached to the bottom of the culture plate, which is substantially lacking cell-to-cell contact, as shown in Figure 1D. Even though in the high-density culture condition, the cell contacts in 2D are only through the cell edges with much loose communication.

**Figure 1.**
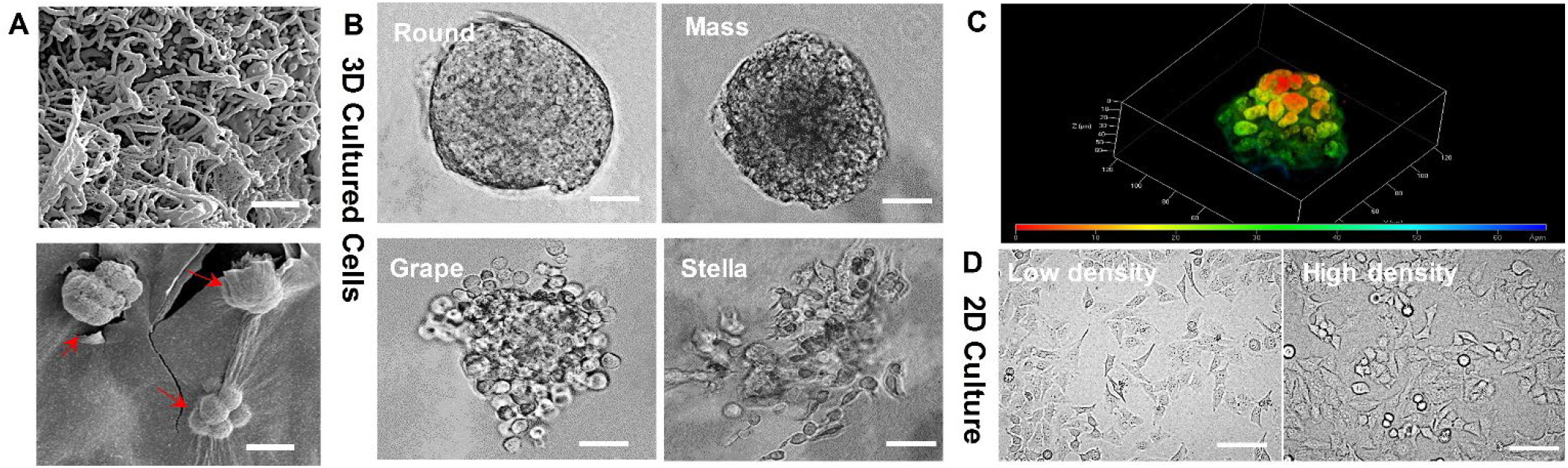
(A) The SEM images showing the morphology of peptide hydrogel as the scaffolds (top, the scale bar is 5μm) for the 3D culture of cervical cancer HeLa cells (bottom, the scale bar is 50 μm). (B) The four typical 3D HeLa cellular morphologies (round, mass, grape, and stella) observed under the bright-field microscopy. The scale bar is 50 μm. (C) The fluorescence confocal microscopic image showing the 3D HeLa tumor-like growth. The 3D confocal image was reconstructed with density distribution using heat map and the depth is 60 μm. (D) The 2D cultured cellular morphology in flat spreading with both low density and high-density seeding, which was observed under the bright-field microscopy. The scale bar is 20 μm.

We further examined the EV secretion rate by using nanoparticle tracking analysis (NTA) to monitor the EVs derived from both 2D culture and 3D culture systems under different culture durations. For 3D cells, the culture can reach 90% confluence after 11 days which is much slower than 2D culture system. So we followed the comparable cellular growth status to collect EVs. For 2D cultured cells, we harvest EVs from media in 6h, 12h, 24h, 36h, and 48h intervals, while 3D culture-derived EVs are harvested at day 5, 7, 9, 11, and 13, individually. The live cells were counted at each time interval for establishing the calibration curve of 3D cell confluency behavior (Figure s2). The EV secretion rate from the 3D cultured cells is more active when cells approach the confluency, which is reflected by three times more EVs secreted after day 9. In contrast, 2D cultured cells reduce the EV secretion and undergo a declining trend, which may attribute to the reduced cell-to-cell contact and communication. The harvested EVs showed typical round, cup shape via SEM imaging in both 2D and 3D culture conditions (Figure 2 B and D), while 3D culture-derived EVs showed the more dense distribution of smaller size. These observations were in agreement with Rocha, et al. that exosomes released by tissue-engineered tumors were smaller than exosomes released by the same cells cultured in monolayer, and the 3D system is more active for producing exosomes^56, 57^. It is interesting that our observation is the same as Rocha, et al. report that the maximum period for conditioned media collection was 48 h for 2D. Rocha, et al. also observed that in 3D this period could be extended for another 4 d without compromising cell viability^56^. We characterized the confluency property for both 2D and 3D cultures (see Figure s2). For reaching to the same confluency as a 2D system (~90%), our HeLa cells in the 3D system require 11 days. However, after 11 days, we observed decreased secretion of EVs at day 13 which may due to the reduced cell viability. Generally, after 90% confluency, cells are required to do subculture to provide more growing space. This observation could still be speculating and more longer-term study will need to give a definitive conclusion. The number of cells used in 3D cultures for producing EVs is considerably lower than that in 2D (seeding density: 3D ~8×10^4^ *vs*. 2D ~5×10^5^). At the end point of harvesting, 3D cells are ~ 9-fold less than 2D cells. The increased secretion of EVs in the 3D system together with Rocha, et al. and other observations ^56, 58^ strengthened the efficiency of the 3D system which could be more useful and accurate for EV functional studies mimicking the in vivo physiological environment.

**Figure 2.**
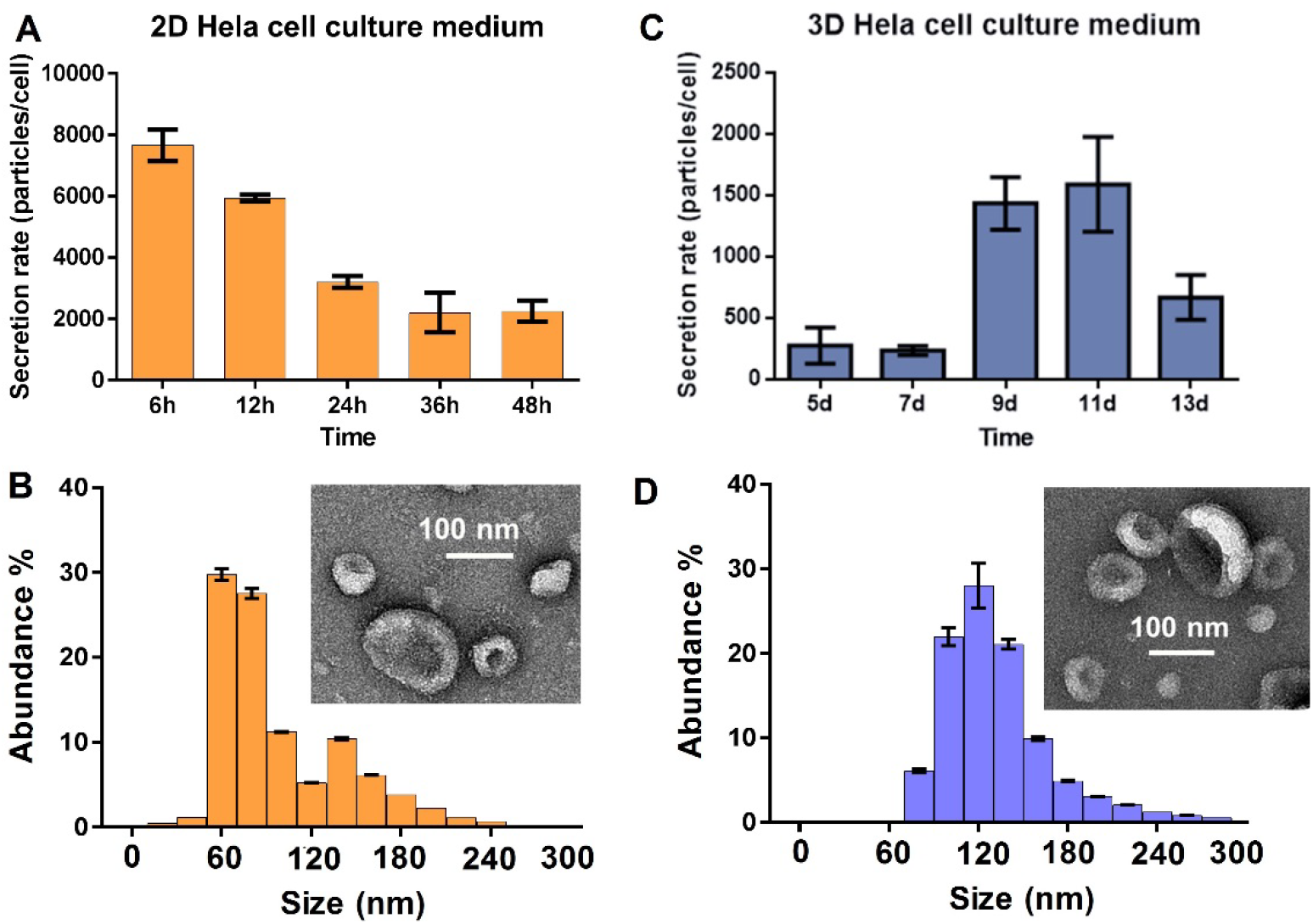
The nanoparticle tracking analysis (NTA) of EV secretion rate between 2D culture (A) and 3D culture (C) systems. The live cell counting was performed at each time interval with three replicated cultures. The statistic CV is ~3-10%. The harvested EVs from 2D (B) and 3D (D) culture conditions showed typical round, cup-shaped EVs around 100 nm size via SEM imaging (Insert). The size distribution and abundance were measured using NTA with five replicated measurements. The statistic CV is ~5%.

### The 3D cell culture derived EV RNA profile shows a significantly high similarity to in vivo EV RNA profile

In order to investigate the EV RNA profiles which are altered under different culture conditions with their parent cells, we prepared multiple EV RNA extracts as analyzed in Figure 3 by Agilent Bioanalyzer, including EV RNAs derived from 2D culture, 3D culture, and cervical cancer patient plasma, compared with parent cell RNAs in 2D and 3D culture conditions. We study two cervical cancer patient plasma samples and one healthy individual plasma sample for comparison with 3D culture derived EVs, because circulating extracellular vesicles and exosomes now are recognized as promising tumor surrogates^59–65^. Circulating EVs can be collected noninvasively with highly increased abundance in cancer patient plasma^64^. Additionally, EVs are relatively stable over time and include proteins and nucleic acids from their parental tumors^66^. Thus, cancer patient plasma could serve as a reference for in vivo tumors^67, 68^.

**Figure 3.**
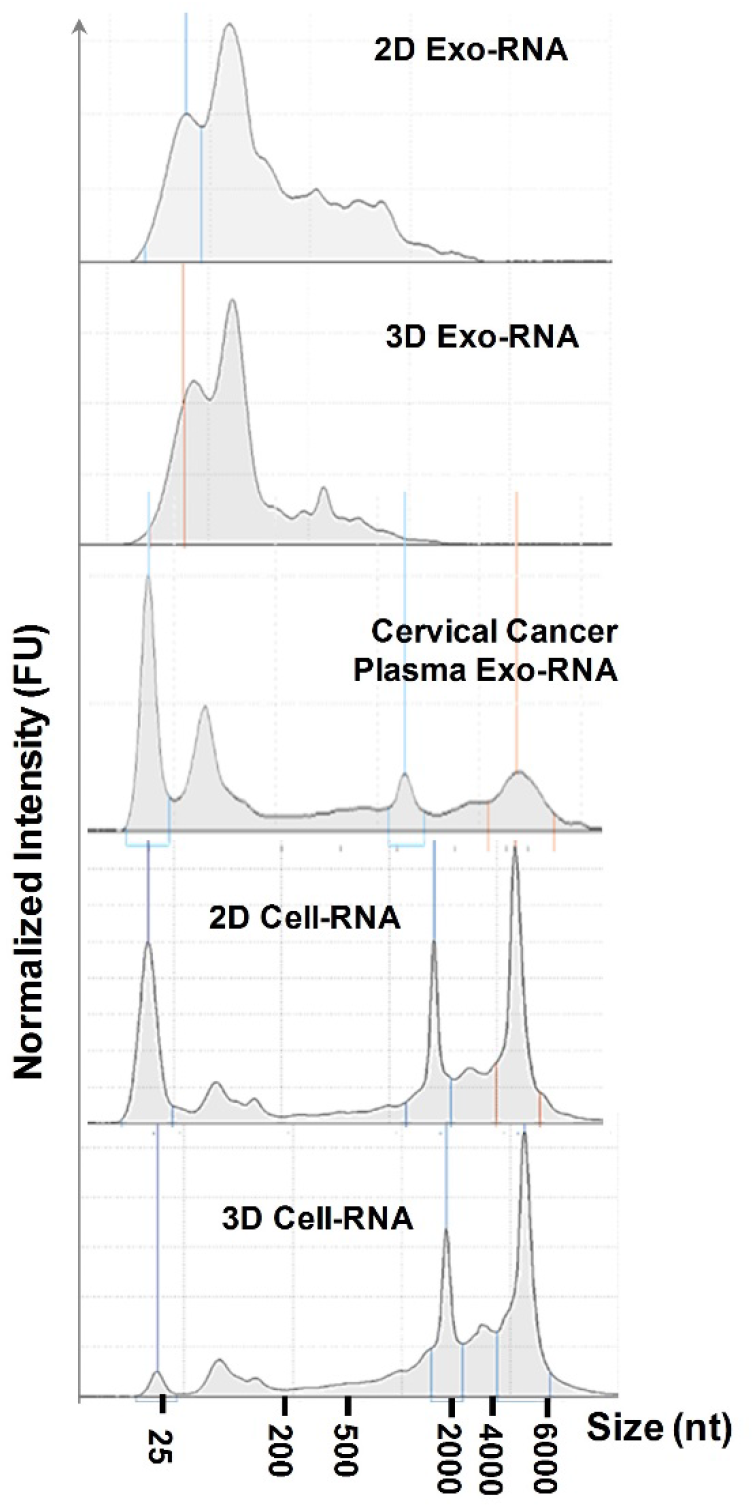
Bioanalyzer analysis of EV RNAs derived from 2D culture, 3D culture, and cervical cancer patient plasma, compared with parent cell RNAs in 2D and 3D culture conditions.

The EV isolation and RNA extraction steps are illustrated in SI Figure s1 and detailed in the experimental section for both cell culture media and human plasma. In contrast to their parent HeLa cell-derived RNAs which are mainly in the long size range (2000-4000 nucleotides), EVs carry more small RNAs around the range below 200 nucleotides. Several studies have shown that EVs carry small coding and noncoding RNAs used by their parent cells as an intercellular network of communication^69–72^. It has been proposed that a selective packaging process for loading RNAs into EVs shedding into extracellular microenvironment which could modify the phenotype of the recipient cells, although the mechanisms underlying are still largely unknown^26, 27^. Parent cells take up solutes, nutrients, and ligands from the extracellular environment via the multiple endocytic pathways and traffic via early endosomes^26^. From endosome maturation into late endosomes, inward budding from the limiting membrane of the endosome leads to the formation of multi-vesicular bodies (MVBs) containing intraluminal vesicles (ILVs), and then fuse with the plasma membrane to release their ILVs into the extracellular space as exosome EVs^26^. During MVB formation, cytosolic RNAs are taken up into ILVs undergoing inward budding with a selective lipid-mediated loading of RNA into EVs^28^. A few RNA-binding proteins have been found capable to selectively bind RNA molecules with specific motifs and induce their export into EVs, such as hnRNPA2B1^26, 73^. In this study, we also can observe a significant amount of small regulatory RNAs enriched in EVs, but not significant in their parent HeLa cells, regardless of 2D or 3D culture conditions. As seen in Figure 3, parent cell-derived RNAs contain a large number of ribosome RNAs in the range of 2000-6000 nucleotides, such as the 16s and 18s. Because the HeLa cell line is the established cell line from the cervical cancer patient, we further compared the EV RNA extract from the cervical cancer patient plasma which also are mainly small coding and noncoding RNAs.

For further understanding the landscape of EV RNA compositions under different culture conditions of their parent cells, and providing an accurate profile of the different coding and non-coding RNA species found per culture compartment, we did small RNAs library preparation (< 200 nt) and used the next-generation sequencing to characterize EV derived RNAs and parent cell RNAs in depth. We generated small RNA libraries from EVs, their donor cells, and corresponding cervical cancer patient plasma samples using NEXTflex small RNA sequencing kit with size selection. The characterization of the small RNA content of EVs is still not well established in this research field and we did substantial optimization work on EV isolation quality control using NTA and SEM/TEM imaging. We also performed the biological replicates for 2D cell RNAs, 3D cell RNAs, and cervical cancer patient plasma EV RNAs. The consistent results between biological replicates indicate that our protocols and results are valid. Thus, the other two biological samples (2D exo-RNAs, 3D exo-RNAs) with one input were considered to be sufficient for all described comparisons. We also used quantitative real-time PCR (qPCR) to validate the NGS results. The resulting reads were aligned to the RefSeq transcriptome annotation (version hg38, downloaded from the UCSC genome browser), the GENCODE (version 28 for hg38), tRNA annotation tracks (version hg38, downloaded from the UCSC genome browser), and the miRNA annotations tracks (version hg38, downloaded from sRNAnalyzer database), using BWA^74^. Only the uniquely aligned reads were considered for downstream analysis. A significant portion (>90%) of the reads were mapped as mRNAs while a much smaller fraction of the reads was mapped as other non-coding RNAs, as shown in Figure 4. This finding is consistent with other reported research that EVs may transport largely mRNA fragments to recipient cells^75, 76^, which may indicate a novel mechanism along with miRNAs for genetic exchange between cells^19^. It is worth mentioning that EV miRNA contents from cervical cancer patient plasma are clearly higher than the healthy individual (3:1), which is consistent with majority reports that miRNAs were significantly enriched in EVs as the important diagnostic markers for cancer^77–79^.

**Figure 4.**
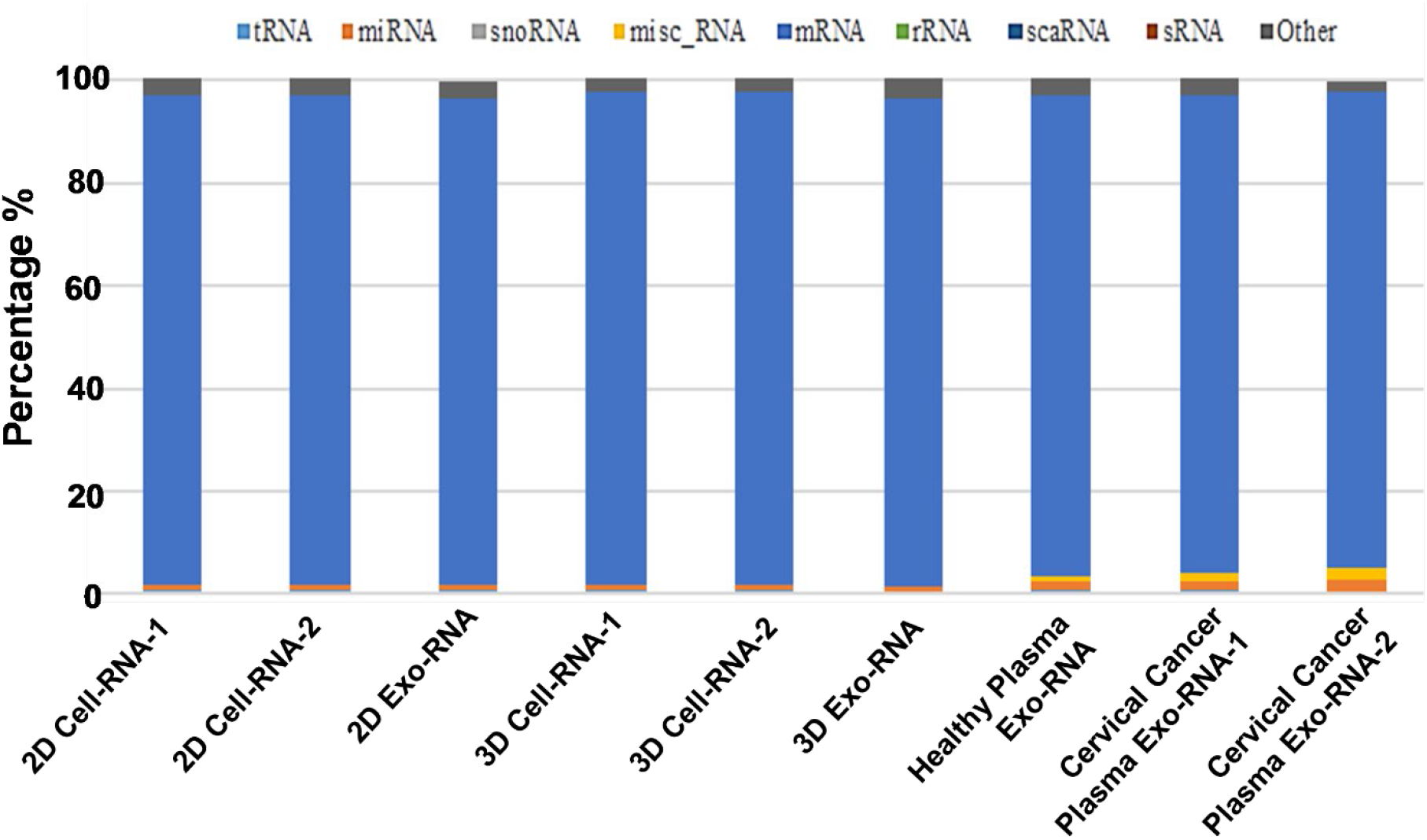
Graph depicting the percentage of reads mapped to transcriptomic regions. The majority of the sequences (>90%) belonged to the protein coding region (mRNA’s) of the transcriptome. The remaining fraction was mapping to the non-coding regions of the transcriptome. Mapping sequences from 2D and 3D cell culture, EVs from 2D and 3D cultures, EVs from cervical cancer patient plasma and EVs from healthy patient plasma cultures were used for analysis. 1 and 2 annotate the two biological replicates.

The next-generation RNA-Seq profiling of EV miRNAs from biopsy specimens is a relatively underexplored frontier. To compare the EV miRNA content derived from various samples, the top 100 miRNAs with the highest average abundances in all samples were selected, and their abundance profiles were subsequently clustered (see Figure 5). EV-derived miRNAs showed significantly different expression profile than their parent cells in both 2D and 3D conditions. However, their parent cells cultured between 2D and 3D systems exhibited similar miRNA expression profile. Rocha, et al. ^56^ also observed that striking differences between small RNAs detected in EVs and in donor cells, which support our results and other research reports indicating a specific sorting process of small RNAs into EVs^26, 27^. As anticipated, replicates (2D-cell miRNAs and 3D-cell miRNAs) were clustered together. It is interesting that we observed the significant difference of miRNAs expression profile between 2D and 3D culture derived EVs, while Rocha, et al showed high similarity^56^. The experimental data explaining this phenomenon are still scarce, and more investigation should be performed for a definitive conclusion. More importantly, we observed that 3D cell-derived EV sample was clustered together with two in vivo cervical cancer patient plasma sample derived EV miRNAs. It supports that 3D cell culture is necessary for reproducing the EV miRNA content sorted by in vivo cells and establishing an accurate disease model^57, 58^. On the other hand, the 2D-derived samples were clustered away from the in vivo samples, indicating that 2D cell culture was unable to represent in vivo biological status. This observation supports the statement from other studies^56, 58^ that 3D culture system would be more useful and accurate for mimicking in vivo physiological environment in studying EV functions. It is interesting that multiple miRNAs were discovered to be present in 3D cell-derived EVs as well as the cervical cancer patient plasma-derived EVs, but not in 2D culture derived EVs. A few of these miRNAs, such as miRNA-2277, miRNA-3691, miRNA-4488, miRNA-548ao, miRNA-6871, miRNA-93, miRNA-4763, miRNA-6751, and miRNA-4633-5p, have been highly associated with control of stability and translation of mRNA ^80 81 82^. To explore the miRNA pathways significantly enriched in EVs from their parent cells, we performed an over-representation analysis (ORA) of two groups of miRNAs with a fold change threshold of 10 (i.e. Log of fold change >10). The results were presented in supplementary information Table s1 and Table s2. The top pathways with 10-fold change or above are mostly associated with metabolism pathway, transport and translation of proteins, and remained similar between EVs and their parents without the influence of culture conditions (2D exo miRNAs *vs*. 2D cell miRNAs, 3D exo miRNAs *vs*. 3D cell miRNAs).

**Figure 5.**
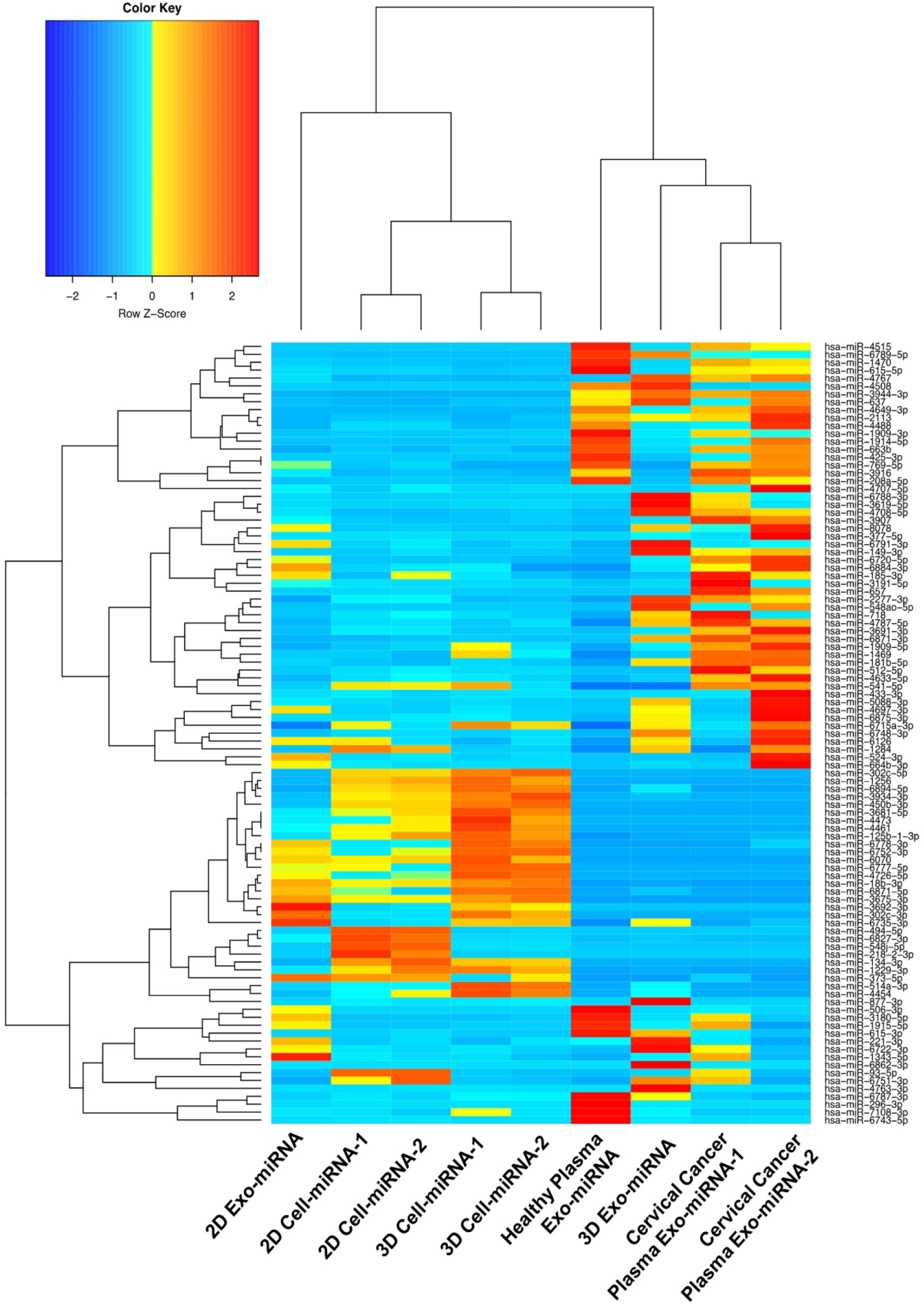
Heatmap with dendrogram depicting top 100 highly expressed miRNAs in 2D Cell, 3D Cell, and their derived EV miRNAs in relation to cervical cancer patient plasma EV miRNAs and healthy plasma EV miRNAs. Red color indicates a higher expression z-score. Hierarchical clustering was performed, using the Spearman correlation method. 3D culture derived EVmiRNA profile and Cervical cancer patient plasma EV miRNA profile have been clustered together due to higher similarities in their transcript expressions. 1 and 2 annotate the two biological replicates.

We further performed region-based annotation on the EV RNA reads, using ANNOVAR^74^. Most of the reads (~90%) belonged to the Intronic, Intergenic, Upstream and UTR5 regions, and their distributions are shown in Figure 6. The remaining reads (10-12%) were mapped to other whole genomic regions (Exonic, Splicing, UTR3, Downstream). The results again showed the significant similarity between the 3D cell-derived EV RNAs and in vivo cervical cancer patient plasma-derived EV RNAs, reconfirming the advantage of using 3D cell culture system.

**Figure 6.**
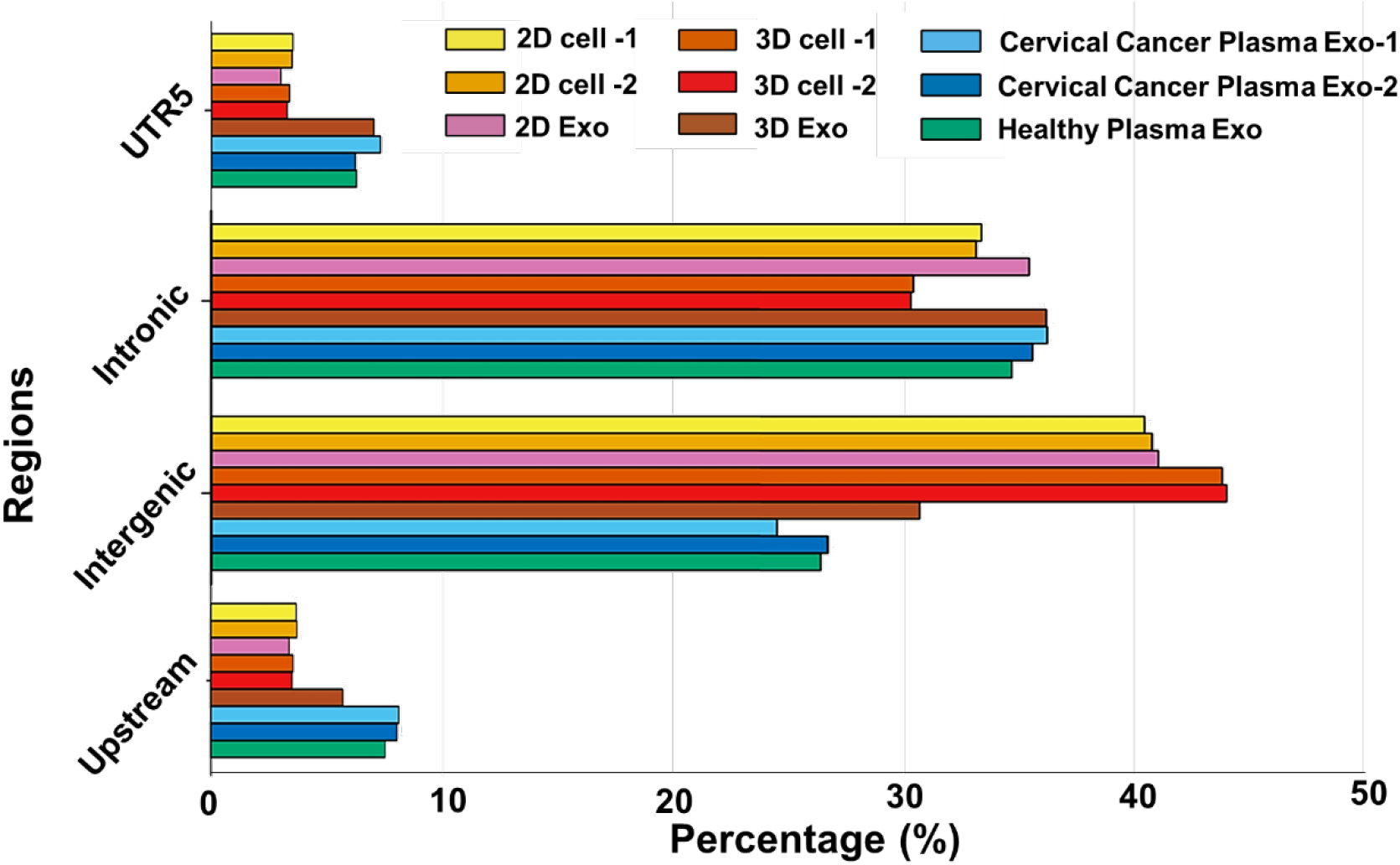
Bar graph depicting the percentage of reads mapped to whole genome regions. The majority of the sequences belong to Intronic, Intergenic, Upstream and UTR5 regions of the whole genome nucleotide sequences. Comparisons were made with nucleotide sequences derived from cervical and healthy patient plasma EVs, 2D cell, 2D EVs, 3D cell, and 3D EVs. 1 and 2 annotate the two biological replicates.

In order to validate the RNA-Seq data and prove the viability of results, we selected six miRNAs from previously samples submitted to RNA-Seq (Hsa-miR-125b-1-3p, Hsa-miR-208a-5p, Hsa-miR-450b-3p, Hsa-miR-1229-3p, Hsa-miR-1284, Hsa-miR-1909-3p) for quantitative real-time PCR validation (Figure s3 and Figure 7). The selected 6 miRNAs must be expressed in all samples (FPKM > 0), with a good sample detection (FPKM > 10,000). We compared expression level fold change between 3D EVs *vs*. 2D EVs, and 3D cells *vs*. 2D cells as shown in the Figure 7. In the cellular system, the expression levels of the six miRNAs mostly are correlated with the data obtained by NGS in terms of the trends of down-regulation or up-regulation, especially for Hsa-miR-125b-1-3p, Hsa-miR-450b-3p, and Hsa-miR-1909-3p which are all upregulated in the similar fold change (~1.5 fold) from both PCR and NGS analysis in 3D culture system. Those three miRNAs are reported to be associated with regulating tumor growth^83–85^. For the EV RNAs data, those three miRNAs (Hsa-miR-125b-1-3p, Hsa-miR-450b-3p, and Hsa-miR-1909-3p) are also correlated very well between PCR and NGS analysis, with much higher fold change (~2-5 fold) in the 3D system. Overall, in the EVs system, the fold change levels are higher than that in the cellular system, which may indicate 3D culture derived EV system carries more enriched miRNAs involved with cellular funcaitonal regulation. No matter 2D culture or 3D culture, among those six miRNAs, Hsa-miR-208a-5p did not show much significant expression level fold change in both EVs system and cellular system under the analysis of both PCR and RNA-Seq. The Hsa-miR-208a-5p is mostly involved in post-transcriptional regulation of gene expression by affecting the stability and translation of mRNAs, which may not be influcenced under different culture conditions. We observed a strong correlation between the quantitative real-time PCR results and the small RNA sequencing data in both cellular RNAs and EV RNAs, and 3D samples showed higher expression levels of the selected miRNAs. This observation also validates our NGS methods for investigating 3D culture system employed in mimicking in vivo tissue system and producing extracellular vesicles.

**Figure 7.**
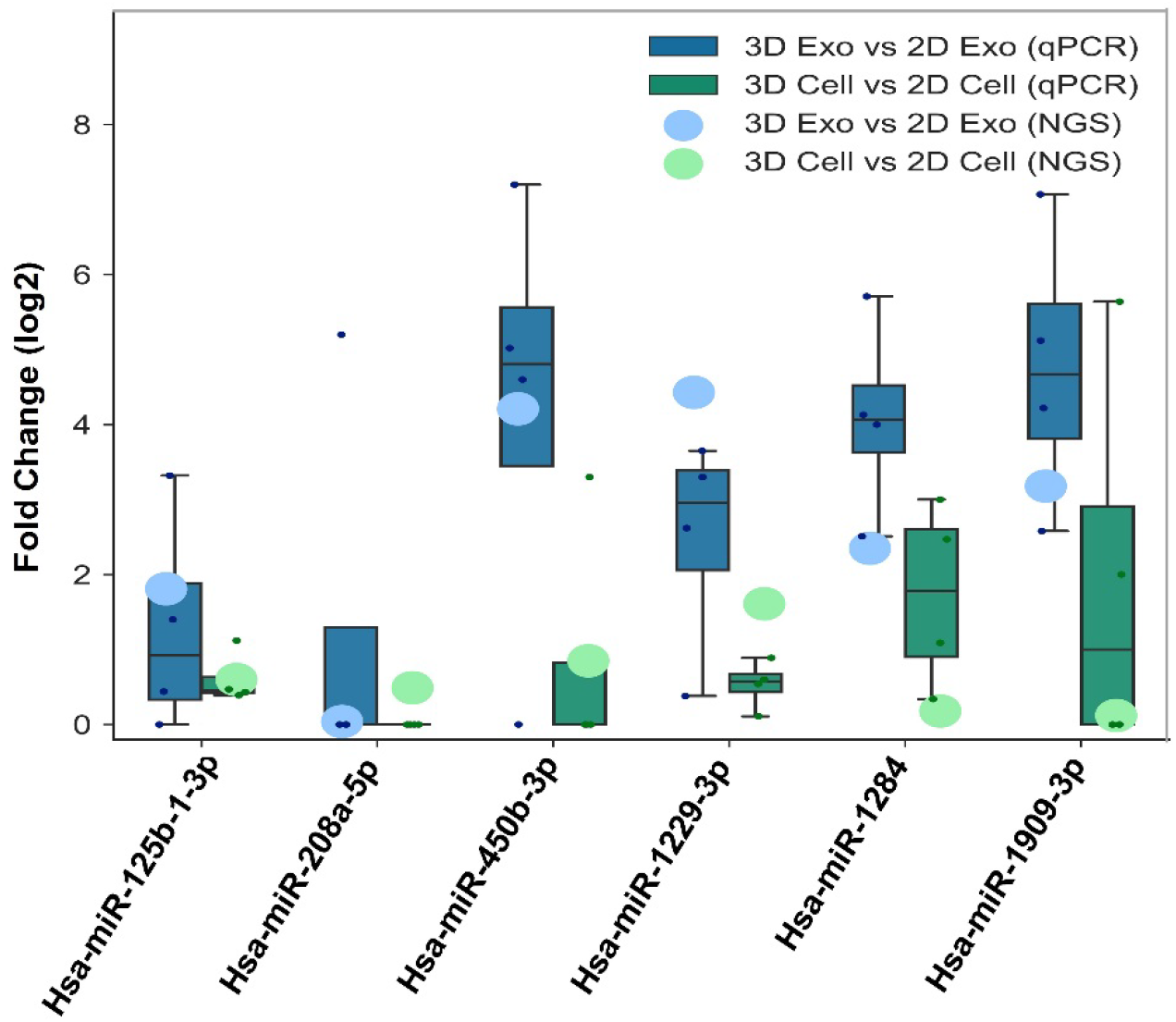
Box-plots depict the correlation between real-time PCR (qPCR) and small RNA Next Generation Sequencing (NGS), for six differentially expressed miRNA genes. Log2-fold changes calculated for both real time qPCR (green and blue boxes) and NGS (green and blue circles) analysis are reported. The lines inside the boxes denote the medians. The boxes mark the interval between the 25th and 75th percentiles with SD as the error bars.

### The 3D cell culture derived EV DNA profiling

We also performed the extraction of EV DNA for analyzing DNA sequences between different cultures of their parent cell. To ensure the quality of extracted EV DNAs without the interference of cell free-DNAs and RNAs, we optimized the extraction protocols using the combination of the QIAamp DNA Mini Kit and the exoRNeasy Serum/Plasma Starter Kit with adding both DNase and RNase sequentially (see experimental section for details). The quality of extracted EV DNA was assessed by Agilent ChipStation is shown in Figure A. Most of the DNA fragments are from genomic DNAs and smaller than 1000 nucleotides. Highly correlated DNA fragment length distributions of the 2D and 3D culturing methods were observed. The reads were mapped to the reference genomes obtained from RefSeq (version hg38), using BWA aligner. The Spearman correlation of the protein-coding gene abundances between samples was calculated and shown in Figure 8B. The results reaffirm that all samples have high abundance correlation, showing that the EV DNA content is more stable than the RNA content, and appears recalcitrant to various culturing methods.

**Figure 8.**
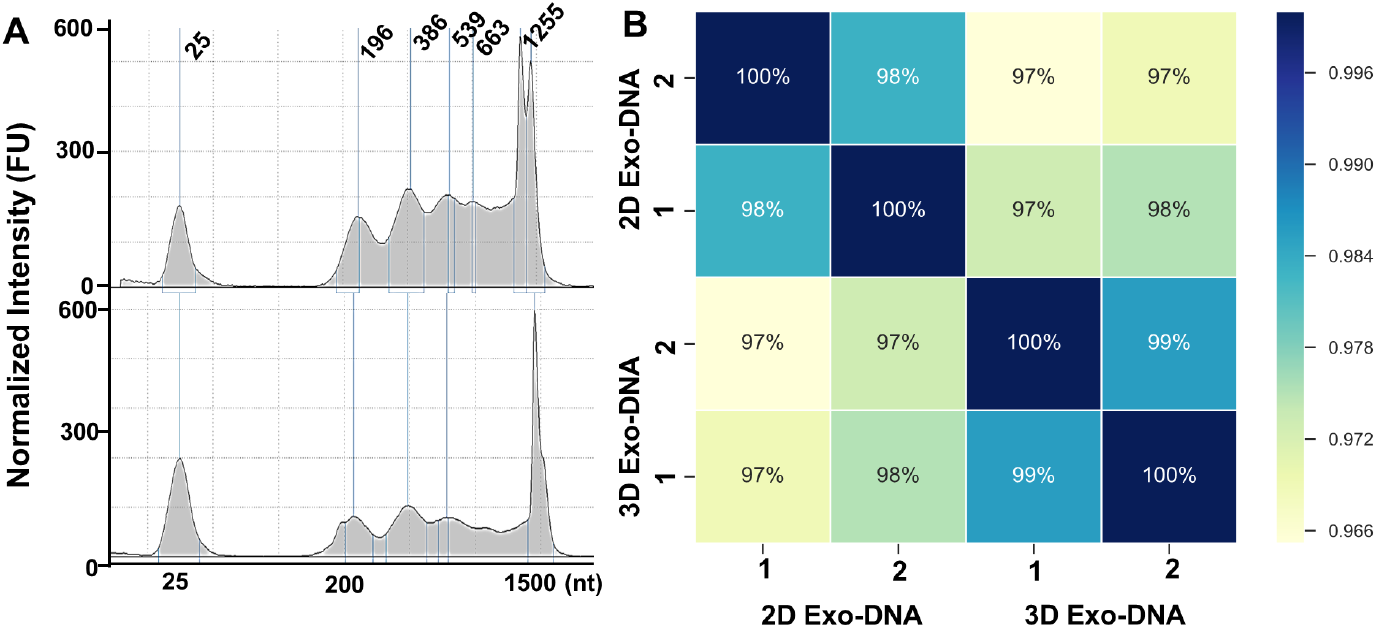
(A) Quality analysis of extracted EV DNAs derived from 2D culture (top) and 3D culture (bottom). (B) The correlation matrix of the DNAseq reads from two biological replicates of 2D and 3D EV DNA samples using the Spearman correlation coefficient. The two biological replicates are exactly consistent with 100% correlation. The 2D and 3D EV DNAs are also highly correlated with similarity coefficient > 95% (2D EV-DNA or 3D EV-DNA). 1 and 2 annotate the two biological replicates.

For the DNA reads that were not mapped as protein-coding genes, we further predict their mapped regions (peaks) using HOMER method^74^. Similar to the protein-coding abundance profile, the predicted peaks also show high abundance correlation between the 2D and 3D samples. Based on the above observations, the culture and growth conditions do not affect the genomic information carried by EV secretion. The recent report^86^ also discovered that serum-derived EV dsDNA was highly consistent with the paired tumor genome, which is essential evidence that EV DNA can be used as a genetic marker for cancer diagnosis via screening gene mutations.

## Discussion

EVs are important non-contact cellular communication pathways for horizontally transferring genetic information. Intensive studies have been devoted to this research field in recent five years showing that EV miRNAs play an important role in disease progression, angiogenesis, and cancer metastasis^87, 88^. The sequencing approaches for studying EV RNAs and DNAs have been a spotlight and not well established in recent two years. However, using the right sample sources and cellular model systems are very critical. We observed that EV-derived miRNAs showed significantly different expression profile than their parent cells in both 2D and 3D conditions. However, their parent cells cultured between 2D and 3D systems exhibited similar miRNA expression profile. More importantly, we observed that 3D cell-derived EV sample was clustered together with two in vivo cervical cancer patient plasma sample derived EV miRNAs. Our study by comparing the secretion dynamics of EVs between 2D culture and 3D culture systems proved that EVs produced in the 3D culture environment may be closer to those produced by patient tumors, which is consistent with recent reports^56, 58^. Thus, 3D culture system may constitute a more useful model for mimicking in vivo physiological environment in studying EV production and functions. For accurately understanding the real cellular communication in a biological system, the in vivo and real-time collected EVs can better reflect the cellular-level communications. However, obtaining human in vivo samples are always challenging due to limited access and regulatory issues. Therefore, building the ex-vivo cellular model is absolutely needed. The 3D culture protocols established in our lab using peptide hydrogel could produce cells with the secretion of in-vivo like EVs. This discovery also can lead to a viable and non-destructive approach for studying tissues and organs via collecting EVs from interstitial fluids or culture medium.

The cellular secretion of EVs is a dynamic process and reflects the microenvironmental changes or parent cell stress. EVs carry miRNAs which can be taken up by neighboring or distant cells for modulating recipient cells. Our observation and other reports^26, 27^ both indicated that miRNA profiles of EVs differ from those of the parent cells, which may due to an active sorting mechanism of EV miRNAs, though this mechanism is still under intensive investigations^26, 27^. Our study compared the EV RNAs derived from 2D culture and 3D culture of cervical cancer cells, in relation to human plasma EV RNAs from cervical cancer patients. This result is the evidence showing that nearly ~ 96% similarity exists between 3D culture derived EV miRNAs and cervical cancer patient plasma-derived EV miRNAs. The region-based annotation on the EV RNA reads also showed that most of the reads (~90%) belonged to the Intronic, Intergenic, Upstream and UTR5 regions from the 3D culture derived EVs are expressed at the same level as the cervical cancer patient plasma-derived EV miRNAs. On the contrary, 2D culture derived EV miRNAs does not correlate as highly with human plasma EV RNA profile (~80%). Our study in this paper indicates that culture conditions could induce the EV RNA change, especially for miRNAs which have been well recognized as the crucial cancer diagnostic biomarkers^89, 90^. By analysis of EV DNA sequencing data, we observed that culture or growth conditions do not affect the genomic information carried by EV secretion, which is supported by recent reports which used EV DNA as their parent tumor surrogate for cancer diagnosis^91, 92^.

We also identified various microRNAs that were common to the 3D culture derived EVs and cervical cancer plasma-derived EVs, indicating a much higher similarity of the 3D culture derived EVs with the cervical cancer plasma-derived EVs, compared to 2D culture derived EVs. These common microRNAs have been shown to be associated with the stability and translation of mRNAs. We plan to submit the RNAs that were identified in our experiments to widely used EV databases like ExoCarta(www.exocarta.org) which is a manually curated database of EV proteins, RNA and lipids. Such databases are excellent sources to obtain information about EV microRNAs, proteins and mRNAs, as provided by other researchers in the field.

Overall, our study focused on establishing a robust 3D cellular system allowing efficient EV production under controlled conditions and mimicking in vivo physiological environment for producing in vivo like EVs, in comparison with conventional 2D cultures. The observation supports our hypothesis that release biogenesis and molecular contents from EVs are very sensitive to the cellular culture environment, which could be an essential evaluation matrix for building biomimetic tissue accurately and employed in drug screening.

## Methods

### 2D and 3D cell culture

Hela cells (ATCC^®^ CCL-2™) were maintained by MEM complete medium (L-glucose MEM medium, Gibco, 11095072) with 10% (v/v) exosome-depleted fetal bovine serum (Gibco, A2720801) at 37 °C with 5% CO_2_. The flask was treated with Poly-D-Lysine Hydrobromide (MP Biomedicals) for reducing variations from substrate interaction. The 1:5 ratio of resuspended cells were taken to a new flask and maintained the culture until the next subculture. To subculture the cell, the cells were digested by 0.25% trypsin (Gibco, 25200-056) and resuspend by fresh MEM medium when the confluence reached to 90%. The culture steps are detailed in IS Figure s1, and the confluence characterization for 2D and 3D culture is described in SI and Figure s2. The 3D culture process was carried out by using the commercial PGmatrix DMEM Kit (PepGel LLC, PGD-006) per vendor’s protocol. Hela cells from the same batch of 2D culture were seeded into the peptide hydrogel with density ~ 8 × 10^4^ which was measured by an automated cell counter (Millipore Corporation, PHCC3000). The 3D culture was maintained using a 24-well cell culture plate with ~ 390 μL cell suspension, 10 μL PGworks solution, and 100 μL PGmatrix solution. The mixture was mixed by pipetting up and down carefully without introducing any bubbles. The well plate was placed into the 37 °C incubators for 1 h. After gelation, 1000 μL warmed fresh MEM complete medium with 10% (v/v) exosome-depleted fetal bovine serum was added to the top of the gel without disturbing. The medium exchange was performed every two days for supplying fresh nutrition and preventing drying out of hydrogel due to long-term culture. The ZOE™ Fluorescent Cell Imager (Bio-Rad Laboratories) was used to observe growth behavior with multiple time intervals.

### Human plasma samples

We used the existing sample resources from the University of Kansas Medical Center /Cancer Center Biospecimen repository which is a public biobank repository, and the data that include individually identifiable private information has not been collected specifically for this work, neither the involvement of human subject. All sample collection methods were carried out in accordance with relevant guidelines and regulations, and there are no specific protocols approval needed.

### EV isolation and NTA analysis

EV isolation steps are illustrated in SI Figure s1. For collecting 2D cultured medium used to characterize EV secretion dynamics, several time intervals were performed according to the confluence behavior (90% confluence in 48 hrs): 6, 12, 24, 36, 48, and 60 hours post subculture. To remove the cell debris, the medium was centrifuged at 3,000g for 15 mins within 4 °C. To collect the supernatant carefully ready for EV isolation, the supernatant was spined at 10,000g for 20 mins, and the debris at the bottom was discarded. For collecting 3D cultured medium used to characterize EV secretion dynamics, below time intervals were performed according to the 3D cell confluence behavior (90% confluency in 11 days): 5, 7, 9, 11, 13 days post subculture. For recovering 3D spheroids, the scaffolding hydrogels were broken mechanically by pipetting up and down, and then transferred into a 15 mL tube for centrifuging at 600 g for 10 min at room temperature for collecting cell pellet. ~0.5 mL 0.25% trypsin (Gibco, 25200-056) was added into the pellet, resuspended gently, and then incubated at 37 °C for 5 min. The trypsin digestion was stopped by adding 0.5 mL MEM complete medium, and the cell pellet was ready for use. For collecting 3D culture derived medium, after breaking scaffolding hydrogels by pipetting up and down, the entire solution was vortexed for 2 minutes and transferred into a 15 mL tube for centrifuging at 600 g for 10 min at room temperature. The supernatant was collected carefully for EV isolation. We use centrifugation protocols developed in our lab to collect EVs-containing supernatant detailed in SI Figure s1. The collected supernatants were subject to a filtration process developed in our lab. The 0.22 μm filter (Millipore Express PLUS (PES) membrane) was conditioned with 10% (v/v) exosome-depleted fetal bovine serum for 2 minutes, then introduce prepared supernatants for filtration. The 10% exo-depleted FBS conditioning could reduce the trap of small EVs when filtration. We did compare the EV particle numbers before and after filtration using NTA analysis and did not see much difference, which is also consistent with the reported filtration method for ensuring the purity of smaller EVs^93^. Afterward, we combine Qiagen ExoRNeasy kit which contains exoEasy spin filtration column. The exoEasy spin column was used for spinning at 500 g for 1min. The flow-through medium was discarded and spin again for 1 min at 3,200 g. ~10 mL XWP buffer was added to the column and spin for 10 min at 3,200 g to wash the trapped exosomes in the membrane of the spin column for following downstream RNA or DNA extraction.

For Nanoparticle Tracking Analysis (NTA), the 2D and 3D media collected at different time intervals follow above-mentioned centrifugation and filtration steps for collecting isolated EVs. The collected solution was used to measure the particle concentration and size distribution using NanoSight LM10 (Malvern Instruments) with standard calibration following the vendor’s instruction.

### EV RNA extraction

The culture media from the 2D and 3D cultures were harvested when the confluence reached ~90% (confluence characterization was shown in Figure s2). Based on the log phase of growth of 3D cells, exosomes were harvested after 11 days of a subculture which is comparable to the confluence as the 2D culture. The medium was prepared for EV isolation based on protocols mentioned above, as well as description in SI Figure s1. The exosomal RNAs were extracted using the exoRNeasy Serum/Plasma Starter Kit (Qiagen, 77023) following the vendor’s instruction. The kit comprises two phases: exosome purification and RNA isolation. The kit uses a spin column format and specialized buffers to purify EVs. Total RNA is then extracted using QIAGEN miRNeasy technology. In the exosome purification stage, above pre-processed medium or human plasma is mixed with Buffer XBP and bound to an exoEasy membrane affinity spin column. The bound exosomes are washed with Buffer XWP, and then lysed with QIAzol. In the RNA extraction step, chloroform is added to the QIAzol eluate, and the aqueous phase is recovered and mixed with ethanol. Total RNA, including miRNA, binds to the spin column, where it is washed three times and eluted. The Agilent Bioanalyzer 2100 Small RNA Chip (Agilent Technologies) was used to determine the quality of the exosomal RNA following the vendor’s instruction by adding 2 μL of extract into Bioanalyzer Chip.

### EV DNA extraction

The EV DNAs were extracted by the combination of the QIAamp DNA Mini Kit (Qiagen, 51304) and the exoRNeasy Serum/Plasma Starter Kit (Qiagen, 77023) following the vendor’s instruction. ~4 mL cell culture media were used to purify EVs. The exoEasy spin column was used for spinning at 500 g for 1min. The flow-through medium was recycled and spin again for 1 min at 3,200 g. ~10 mL XWP buffer was added to the column and spin for 10 min at 3,200 g to wash the trapped EVs on the membrane of the spin column. The QIAamp DNA Mini Kit was used to extract the EV DNAs. ~ 400 μL PBS buffer, 40 μL QIAGEN Protease, and 4 μL RNase (Qiagen, 19101) were mixed into one 1.5 microcentrifuge tube and transferred into the center of the column membrane for wetting the membrane. ~400 μL AL buffer was added to the membrane slowly and mixed by pipette up and down without touching the membrane. The resulted column was incubated for 10 min at 56 °C, and then spin for 10 min at 3,200 g to collect the flow-through lysate into a new 1.5 mL microcentrifuge tube. ~ 400 μL ethanol was added to the lysate and mixed for 15 s. The QIAmp Mini spin column was used for spinning at 6,000g for 1 minute. Then the column membrane was washed with 500 μL AW1 buffer and AW2 buffer separately provided in the kit. The DNA concentration was measured by Qubit^®^ 3.0 Fluorometer (Life Technologies, Q33216).

### Sequencing and data analysis

The RNA library was prepared by using the commercial library preparation kit NEXTflex small RNA sequencing kit (Bioo Scientific, NOVA-5132-05) following the recommended protocol by the manufactures. About ~20 ng input quantities was used with this kit. PAGE-based size selection is used for handling total extracted RNAs for 25 cycles of PCR. The Next-generation sequencing libraries for extracted EV DNAs was prepared using Illumina Nextera XT DNA Library Preparation Kit. The DNA peaks with 150 bp were purified by the gel size selection approach and eluted in ~ 8 μL elution buffer. The both amplified libraries (RNAs and DNAs) were examined by the TapeStation gel (Angilent 2200) and quantified by Qubit^®^ 3.0 Fluorometer. The sequencing was performed by the KU Genomic Core using the Illumina HiSeq 2500 System with about ~3 million reads per sample. Raw reads were trimmed to remove the 3’ adaptor sequences. The sequences were filtered by removing sequences with lengths <16 nt. The filtered sequences and quantified by were mapped onto the RefSeq transcriptome annotation (version hg38, downloaded from the UCSC genome browser), GENCODE (version 28 for hg38), tRNA annotation tracks (version hg38, downloaded from the UCSC genome browser), and the miRNA annotations tracks (version hg38, downloaded from sRNAnalyzer database). The RefSeq whole genome annotation was used to perform region-based annotation. Mapping was performed using BWA Aligner. Only a single optimal hit was retrieved, by setting the parameter R=1, during alignment. Differences in seed length were disallowed (set k=0). All other parameters were set to default. A comparison between the percentage of reads mapped using a single hit parameter (R=1) and the default parameter, which allows multiple hits (R=30) is shown in Table 1. It was observed that the mapping percentages were higher when multiple hits were allowed, as opposed to a single best hit. However, since the sample sequences used were very short (<100bps), the sub-optimal hits obtained were most likely to be false positive alignments, thereby making them unsuitable for our analysis purposes.

**Table 1.**
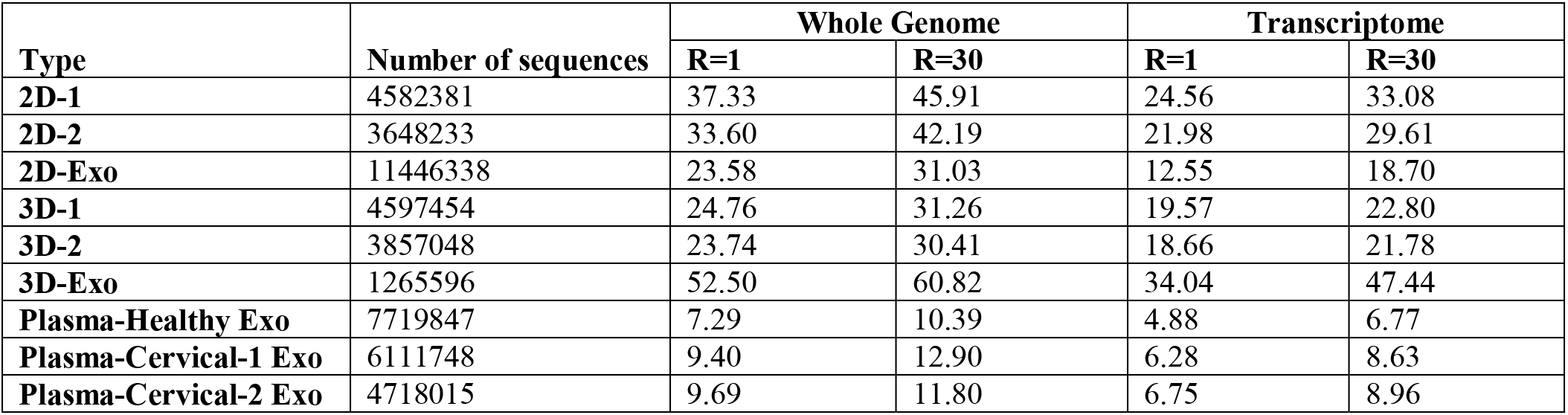
BWA mapping percentages of filtered nucleotide sequences with parameters –R 1 (only single hit) and –R 30 (default parameter-allows multiple hits for whole genomic and transcriptomic regions.

| Type | Number of sequences | Whole Genome |  | Transcriptome |  |
| --- | --- | --- | --- | --- | --- |
|  |  | R=1 | R=30 | R=1 | R=30 |
| 2D-1 | 4582381 | 37.33 | 45.91 | 24.56 | 33.08 |
| 2D-2 | 3648233 | 33.60 | 42.19 | 21.98 | 29.61 |
| 2D-Exo | 11446338 | 23.58 | 31.03 | 12.55 | 18.70 |
| 3D-1 | 4597454 | 24.76 | 31.26 | 19.57 | 22.80 |
| 3D-2 | 3857048 | 23.74 | 30.41 | 18.66 | 21.78 |
| 3D-Exo | 1265596 | 52.50 | 60.82 | 34.04 | 47.44 |
| Plasma-Healthy Exo | 7719847 | 7.29 | 10.39 | 4.88 | 6.77 |
| Plasma-Cervical-1 Exo | 6111748 | 9.40 | 12.90 | 6.28 | 8.63 |
| Plasma-Cervical-2 Exo | 4718015 | 9.69 | 11.80 | 6.75 | 8.96 |

### Quantitative real-time PCR

**For validating small RNA sequencing data and prove the viability of the results**, TaqMan Advanced microRNA assays (Thermo Fisher Scientific) were used to quantify specific microRNAs from parent cells and derived EV RNA samples, on a 7500 Fast Real-Time PCR system (Thermo Fisher Scientific). The samples previously submitted to small RNA sequencing were used and we selected 6 small RNAs that must be expressed in all samples (FPKM > 0), with a good sample detection (FPKM > 10,000), and compared the fold change between two groups of samples. Briefly, 100 ng of RNA was reversed transcribed using universal RT primers and the TaqMan Advanced microRNA cDNA Synthesis Kit (Thermo Fisher Scientific), according to vendor’s instructions. Afterwards, real-time PCR was performed for each sample using 10 μL of diluted cDNA, TaqMan Advanced microRNA assays, and TaqMan Fast Advanced Master Mix (Thermo Fisher Scientific). Reactions were done with four repeats and data were analyzed by the comparative 2^ΔCT^ method, where ΔCT = C(t)miRNA 3D cells - C(t)miRNA 2D cells or ΔCT = C(t)miRNA 3D EVs - C(t)miRNA 2D EVs. Results were represented as mean ± standard deviation. The detailed microRNA assays and PCR analysis were shown in Figure s3 in the Supplemental Information.

## Acknowledgments

We thank Lanjing Wei for preparing cell culture derived exosomes, as well as the isolation of human plasma-derived exosomes, and extraction of RNAs and DNAs. We thank Dr. Nan He for performing the quantitative real-time PCR for validating NGS data. We thank Dr. Qingfu Zhu for helping with the preparation of 3D cell culture and TEM imaging of 3D cultured cells. We also thank John Sibbitts for developing confocal imaging of 3D cells and developing exosomal RNA/DNA extraction protocols. We acknowledge the funding support from USDA-NIFA KS451214 and NIH NIGMS P20 GM103638. We also thank the assistant from Jennifer Hackett from the Genomic Core at the University of Kansas for library preparation, quality check and next-generation sequencing. We acknowledge the support of The University of Kansas Cancer Center’s Biospecimen Repository Core Facility staff, funded in part by the National Cancer Institute Cancer Center Support Grant P30 CA168524.

## Author contributions statement

Miss Sirisha Thippabhotla drafted the manuscript, performed database searching, bioinformatics analysis and statistical data analysis, and draw the data figures. Dr. Cuncong Zhong performed overall data collection and analysis, as well as the manuscript revision. Dr. Mei He conceived the research idea, analyzed the data, and revised the manuscript.

## Additional information

There is no conflict of interest to disclose.

The sequence data is made available from exosomal databases ExoCarta or upon request.

## Competing interests

The corresponding author is responsible for submitting a competing interests statement on behalf of all authors of the paper. The author(s) declare no competing interests.

## References

1. van Niel, G., D’Angelo, G. & Raposo, G. Shedding light on the cell biology of extracellular vesicles. Nat Rev Mol Cell Biol 19, 213–228 (2018).

2. Hessvik, N.P. & Llorente, A. Current knowledge on exosome biogenesis and release. Cell Mol Life Sci 75, 193–208 (2018).

3. Shen, B., Fang, Y., Wu, N. & Gould, S.J. Biogenesis of the posterior pole is mediated by the exosome/microvesicle protein-sorting pathway. J Biol Chem 286, 44162–44176 (2011).

4. Pant, S., Hilton, H. & Burczynski, M.E. The multifaceted exosome: biogenesis, role in normal and aberrant cellular function, and frontiers for pharmacological and biomarker opportunities. Biochem Pharmacol 83, 1484–1494 (2012).

5. Fares, J., Kashyap, R. & Zimmermann, P. Syntenin: Key player in cancer exosome biogenesis and uptake? Cell Adh Migr 11, 124–126 (2017).

6. Raposo, G. & Stoorvogel, W. Extracellular vesicles: Exosomes, microvesicles, and friends. The Journal of Cell Biology 200, 373–383 (2013).

7. Shao, H. et al. New Technologies for Analysis of Extracellular Vesicles. Chem Rev 118, 1917–1950 (2018).

8. Kruger, S. et al. Molecular characterization of exosome-like vesicles from breast cancer cells. BMC Cancer 14, 44 (2014).

9. Saeedi Borujeni, M.J. et al. Molecular aspects of diabetes mellitus: Resistin, microRNA, and exosome. J Cell Biochem 119, 1257–1272 (2018).

10. Thery, C. et al. Minimal information for studies of extracellular vesicles 2018 (MISEV2018): a position statement of the International Society for Extracellular Vesicles and update of the MISEV2014 guidelines. J Extracell Vesicles 7, 1535750 (2018).

11. Wan, Z. et al. Exosome-mediated cell-cell communication in tumor progression. Am J Cancer Res 8, 1661–1673 (2018).

12. Samuelson, I. & Vidal-Puig, A.J. Fed-EXosome: extracellular vesicles and cell-cell communication in metabolic regulation. Essays Biochem 62, 165–175 (2018).

13. Maia, J., Caja, S., Strano Moraes, M.C., Couto, N. & Costa-Silva, B. Exosome-Based Cell-Cell Communication in the Tumor Microenvironment. Front Cell Dev Biol 6, 18 (2018).

14. Zhang, W., Peng, P. & Shen, K. Role of Exosome Shuttle RNA in Cell-to-Cell Communication. Zhongguo Yi Xue Ke Xue Yuan Xue Bao 38, 480–483 (2016).

15. Zhang, D. et al. Exosome-Mediated Small RNA Delivery: A Novel Therapeutic Approach for Inflammatory Lung Responses. Mol Ther 26, 2119–2130 (2018).

16. Jia, Y. et al. Exosome: emerging biomarker in breast cancer. Oncotarget 8, 41717–41733 (2017).

17. Skog, J. et al. Glioblastoma microvesicles transport RNA and protein that promote tumor growth and provide diagnostic biomarkers. Nature cell biology 10, 1470–1476 (2008).

18. Shen, J. et al. Advances of exosome in the development of ovarian cancer and its diagnostic and therapeutic prospect. Onco Targets Ther 11, 2831–2841 (2018).

19. Valadi, H. et al. Exosome-mediated transfer of mRNAs and microRNAs is a novel mechanism of genetic exchange between cells. Nat Cell Biol 9, 654–659 (2007).

20. Wang, J. & Barr, M.M. Cell-cell communication via ciliary extracellular vesicles: clues from model systems. Essays Biochem 62, 205–213 (2018).

21. Mathieu, M., Martin-Jaular, L., Lavieu, G. & Thery, C. Specificities of secretion and uptake of exosomes and other extracellular vesicles for cell-to-cell communication. Nat Cell Biol 21, 9–17 (2019).

22. Hwang, I. Cell-cell communication via extracellular membrane vesicles and its role in the immune response. Mol Cells 36, 105–111 (2013).

23. Tetta, C., Ghigo, E., Silengo, L., Deregibus, M.C. & Camussi, G. Extracellular vesicles as an emerging mechanism of cell-to-cell communication. Endocrine 44, 11–19 (2013).

24. Turturici, G., Tinnirello, R., Sconzo, G. & Geraci, F. Extracellular membrane vesicles as a mechanism of cell-to-cell communication: advantages and disadvantages. Am J Physiol Cell Physiol 306, C621–633 (2014).

25. Wendler, F., Stamp, G.W. & Giamas, G. Tumor-Stromal Cell Communication: Small Vesicles Signal Big Changes. Trends Cancer 2, 326–329 (2016).

26. Zhang, J. et al. Exosome and exosomal microRNA: trafficking, sorting, and function. Genomics Proteomics Bioinformatics 13, 17–24 (2015).

27. Villarroya-Beltri, C., Baixauli, F., Gutierrez-Vazquez, C., Sanchez-Madrid, F. & Mittelbrunn, M. Sorting it out: regulation of exosome loading. Semin Cancer Biol 28, 3–13 (2014).

28. Subra, C., Laulagnier, K., Perret, B. & Record, M. Exosome lipidomics unravels lipid sorting at the level of multivesicular bodies. Biochimie 89, 205–212 (2007).

29. Lampe, K.J., Mooney, R.G., Bjugstad, K.B. & Mahoney, M.J. Effect of macromer weight percent on neural cell growth in 2D and 3D nondegradable PEG hydrogel culture. J Biomed Mater Res A 94, 1162–1171 (2010).

30. Tibbitt, M.W. & Anseth, K.S. Hydrogels as Extracellular Matrix Mimics for 3D Cell Culture. Biotechnology and bioengineering 103, 655–663 (2009).

31. Honegger, A. et al. Dependence of Intracellular and Exosomal microRNAs on Viral E6/E7 Oncogene Expression in HPV-positive Tumor Cells. PLoS Pathogens 11, e1004712 (2015).

32. Kahlert, C. et al. Identification of Double-stranded Genomic DNA Spanning All Chromosomes with Mutated KRAS and p53 DNA in the Serum Exosomes of Patients with Pancreatic Cancer. The Journal of Biological Chemistry 289, 3869–3875 (2014).

33. Taylor, D.D. & Gercel-Taylor, C. MicroRNA signatures of tumor-derived exosomes as diagnostic biomarkers of ovarian cancer. Gynecologic oncology 110, 13–21 (2008).

34. Zhao, Z., Yang, Y., Zeng, Y. & He, M. A Microfluidic ExoSearch Chip for Multiplexed Exosome Detection Towards Blood-based Ovarian Cancer Diagnosis(). Lab on a chip 16, 489–496 (2016).

35. Pampaloni, F., Reynaud, E.G. & Stelzer, E.H.K. The third dimension bridges the gap between cell culture and live tissue. Nature Reviews Molecular Cell Biology 8, 839 (2007).

36. Li, Y. & Kilian, K.A. Bridging the Gap: From 2D Cell Culture to 3D Microengineered Extracellular Matrices. Adv Healthc Mater 4, 2780–2796 (2015).

37. Marino, S., Bishop, R.T., de Ridder, D., Delgado-Calle, J. & Reagan, M.R. 2D and 3D In Vitro Co-Culture for Cancer and Bone Cell Interaction Studies. Methods Mol Biol 1914, 71–98 (2019).

38. Duval, K. et al. Modeling Physiological Events in 2D vs. 3D Cell Culture. Physiology (Bethesda) 32, 266–277 (2017).

39. Lou, E., O’Hare, P., Subramanian, S. & Steer, C.J. Lost in translation: applying 2D intercellular communication via tunneling nanotubes in cell culture to physiologically relevant 3D microenvironments. FEBS J 284, 699–707 (2017).

40. Barros, A.S., Costa, E.C., Nunes, A.S., de Melo-Diogo, D. & Correia, I.J. Comparative study of the therapeutic effect of Doxorubicin and Resveratrol combination on 2D and 3D (spheroids) cell culture models. Int J Pharm 551, 76–83 (2018).

41. Souza, A.G. et al. Comparative Assay of 2D and 3D Cell Culture Models: Proliferation, Gene Expression and Anticancer Drug Response. Curr Pharm Des 24, 1689–1694 (2018).

42. Breslin, S. & O’Driscoll, L. Three-dimensional cell culture: the missing link in drug discovery. Drug Discovery Today 18, 240–249 (2013).

43. Sun, T., Jackson, S., Haycock, J.W. & MacNeil, S. Culture of skin cells in 3D rather than 2D improves their ability to survive exposure to cytotoxic agents. Journal of Biotechnology 122, 372–381 (2006).

44. Kleinman, H.K., Philp, D. & Hoffman, M.P. Role of the extracellular matrix in morphogenesis. Current Opinion in Biotechnology 14, 526–532 (2003).

45. Katsu, M. et al. MicroRNA expression profiles of neuron-derived extracellular vesicles in plasma from patients with amyotrophic lateral sclerosis. Neurosci Lett 708, 134176 (2019).

46. Takov, K., Yellon, D.M. & Davidson, S.M. Comparison of small extracellular vesicles isolated from plasma by ultracentrifugation or size-exclusion chromatography: yield, purity and functional potential. J Extracell Vesicles 8, 1560809 (2019).

47. Minciacchi, V.R., Freeman, M.R. & Di Vizio, D. Extracellular Vesicles in Cancer: Exosomes, Microvesicles and the Emerging Role of Large Oncosomes. Seminars in cell & developmental biology 40, 41–51 (2015).

48. Chatterjee, K., Young, M.F. & Simon, C.G., Jr. Fabricating gradient hydrogel scaffolds for 3D cell culture. Comb Chem High Throughput Screen 14, 227–236 (2011).

49. Benson, K., Galla, H.J. & Kehr, N.S. Cell adhesion behavior in 3D hydrogel scaffolds functionalized with D- or L-aminoacids. Macromol Biosci 14, 793–798 (2014).

50. Gu, J., Zhao, Y., Guan, Y. & Zhang, Y. Effect of particle size in a colloidal hydrogel scaffold for 3D cell culture. Colloids Surf B Biointerfaces 136, 1139–1147 (2015).

51. Wang, X. et al. Self-assembling peptide hydrogel scaffolds support stem cell-based hair follicle regeneration. Nanomedicine 12, 2115–2125 (2016).

52. Reis, L.A. et al. A peptide-modified chitosan-collagen hydrogel for cardiac cell culture and delivery. Acta Biomater 8, 1022–1036 (2012).

53. Zhao, Y. et al. Three-dimensional printing of Hela cells for cervical tumor model in vitro. Biofabrication 6, 035001 (2014).

54. Kenny, P.A. et al. The morphologies of breast cancer cell lines in three-dimensional assays correlate with their profiles of gene expression. Mol Oncol 1, 84–96 (2007).

55. Sant, S. & Johnston, P.A. The production of 3D tumor spheroids for cancer drug discovery. Drug Discov Today Technol 23, 27–36 (2017).

56. Rocha, S. et al. 3D Cellular Architecture Affects MicroRNA and Protein Cargo of Extracellular Vesicles. Adv Sci (Weinh) 6, 1800948 (2019).

57. Villasante, A. et al. Recapitulating the Size and Cargo of Tumor Exosomes in a Tissue-Engineered Model. Theranostics 6, 1119–1130 (2016).

58. Eguchi, T. et al. Organoids with cancer stem cell-like properties secrete exosomes and HSP90 in a 3D nanoenvironment. PLoS One 13, e0191109 (2018).

59. Whiteside, T.L. The emerging role of plasma exosomes in diagnosis, prognosis and therapies of patients with cancer. Contemp Oncol (Pozn) 22, 38–40 (2018).

60. He, M. & Zeng, Y. Microfluidic Exosome Analysis toward Liquid Biopsy for Cancer. J Lab Autom 21, 599–608 (2016).

61. Giannopoulou, L., Zavridou, M., Kasimir-Bauer, S. & Lianidou, E.S. Liquid biopsy in ovarian cancer: the potential of circulating miRNAs and exosomes. Transl Res 205, 77–91 (2019).

62. Halvaei, S. et al. Exosomes in Cancer Liquid Biopsy: A Focus on Breast Cancer. Mol Ther Nucleic Acids 10, 131–141 (2018).

63. Zhang, P., Samuel, G., Crow, J., Godwin, A.K. & Zeng, Y. Molecular assessment of circulating exosomes toward liquid biopsy diagnosis of Ewing sarcoma family of tumors. Transl Res 201, 136–153 (2018).

64. Zhang, W. et al. Liquid Biopsy for Cancer: Circulating Tumor Cells, Circulating Free DNA or Exosomes? Cell Physiol Biochem 41, 755–768 (2017).

65. Zhao, Y. et al. Liquid Biopsy of Vitreous Reveals an Abundant Vesicle Population Consistent With the Size and Morphology of Exosomes. Transl Vis Sci Technol 7, 6 (2018).

66. Eguchi, A., Kostallari, E., Feldstein, A.E. & Shah, V.H. Extracellular vesicles, the liquid biopsy of the future. J Hepatol (2019).

67. Srivastava, A. et al. A Non-invasive Liquid Biopsy Screening of Urine-Derived Exosomes for miRNAs as Biomarkers in Endometrial Cancer Patients. AAPS J 20, 82 (2018).

68. Yoshioka, Y., Katsuda, T. & Ochiya, T. Extracellular vesicles and encapusulated miRNAs as emerging cancer biomarkers for novel liquid biopsy. Jpn J Clin Oncol 48, 869–876 (2018).

69. Lin, Z. et al. Selective enrichment of microRNAs in extracellular matrix vesicles produced by growth plate chondrocytes. Bone 88, 47–55 (2016).

70. Ahadi, A., Khoury, S., Losseva, M. & Tran, N. A comparative analysis of lncRNAs in prostate cancer exosomes and their parental cell lines. Genom Data 9, 7–9 (2016).

71. Hessvik, N.P., Phuyal, S., Brech, A., Sandvig, K. & Llorente, A. Profiling of microRNAs in exosomes released from PC-3 prostate cancer cells. Biochim Biophys Acta 1819, 1154–1163 (2012).

72. Perez-Boza, J., Lion, M. & Struman, I. Exploring the RNA landscape of endothelial exosomes. RNA 24, 423–435 (2018).

73. Tian, T. et al. Dynamics of exosome internalization and trafficking. J Cell Physiol 228, 1487–1495 (2013).

74. Zhong, C. & Zhang, S. Accurate and Efficient Mapping of the Cross-Linked microRNA-mRNA Duplex Reads. iScience 18, 11–19 (2019).

75. Batagov, A.O. & Kurochkin, I.V. Exosomes secreted by human cells transport largely mRNA fragments that are enriched in the 3’-untranslated regions. Biol Direct 8, 12 (2013).

76. Wei, Z. et al. Coding and noncoding landscape of extracellular RNA released by human glioma stem cells. Nat Commun 8, 1145 (2017).

77. Sun, Z. et al. Effect of exosomal miRNA on cancer biology and clinical applications. Mol Cancer 17, 147 (2018).

78. Bhome, R. et al. Exosomal microRNAs (exomiRs): Small molecules with a big role in cancer. Cancer Lett 420, 228–235 (2018).

79. Chen, M. et al. Distinct shed microvesicle and exosome microRNA signatures reveal diagnostic markers for colorectal cancer. PLoS One 14, e0210003 (2019).

80. Kahraman, M. et al. Technical Stability and Biological Variability in MicroRNAs from Dried Blood Spots: A Lung Cancer Therapy-Monitoring Showcase. Clin Chem 63, 1476–1488 (2017).

81. Slattery, M.L. et al. Infrequently expressed miRNAs in colorectal cancer tissue and tumor molecular phenotype. Mod Pathol 31, 209 (2018).

82. Huang, X. et al. Characterization of human plasma-derived exosomal RNAs by deep sequencing. BMC Genomics 14, 319 (2013).

83. Sun, Y.M., Lin, K.Y. & Chen, Y.Q. Diverse functions of miR-125 family in different cell contexts. J Hematol Oncol 6, 6 (2013).

84. Chen, S.F. et al. Identification of core aberrantly expressed microRNAs in serous ovarian carcinoma. Oncotarget 9, 20451–20466 (2018).

85. Duellman, T., Warren, C. & Yang, J. Single nucleotide polymorphism-specific regulation of matrix metalloproteinase-9 by multiple miRNAs targeting the coding exon. Nucleic Acids Res 42, 5518–5531 (2014).

86. Wang, L. et al. Exosomal double-stranded DNA as a biomarker for the diagnosis and preoperative assessment of pheochromocytoma and paraganglioma. Mol Cancer 17, 28 (2018).

87. Kyuno, D., Zhao, K., Bauer, N., Ryschich, E. & Zoller, M. Therapeutic Targeting Cancer-Initiating Cell Markers by Exosome miRNA: Efficacy and Functional Consequences Exemplified for claudin7 and EpCAM. Transl Oncol 12, 191–199 (2019).

88. Manna, I. et al. Exosome-associated miRNA profile as a prognostic tool for therapy response monitoring in multiple sclerosis patients. FASEB J 32, 4241–4246 (2018).

89. Valencia, K. et al. miRNA cargo within exosome-like vesicle transfer influences metastatic bone colonization. Mol Oncol 8, 689–703 (2014).

90. Madhavan, B. et al. Combined evaluation of a panel of protein and miRNA serum-exosome biomarkers for pancreatic cancer diagnosis increases sensitivity and specificity. Int J Cancer 136, 2616–2627 (2015).

91. Iliev, D. et al. Stimulation of exosome release by extracellular DNA is conserved across multiple cell types. FEBS J (2018).

92. Smith, R.A. & Lam, A.K. Liquid Biopsy for Investigation of Cancer DNA in Esophageal Adenocarcinoma: Cell-Free Plasma DNA and Exosome-Associated DNA. Methods Mol Biol 1756, 187–194 (2018).

93. Gheinani, A.H. et al. Improved isolation strategies to increase the yield and purity of human urinary exosomes for biomarker discovery. Sci Rep 8, 3945 (2018).

