## Supplemental Files for "3D cell culture stimulates the secretion of in vivo like extracellular vesicles"

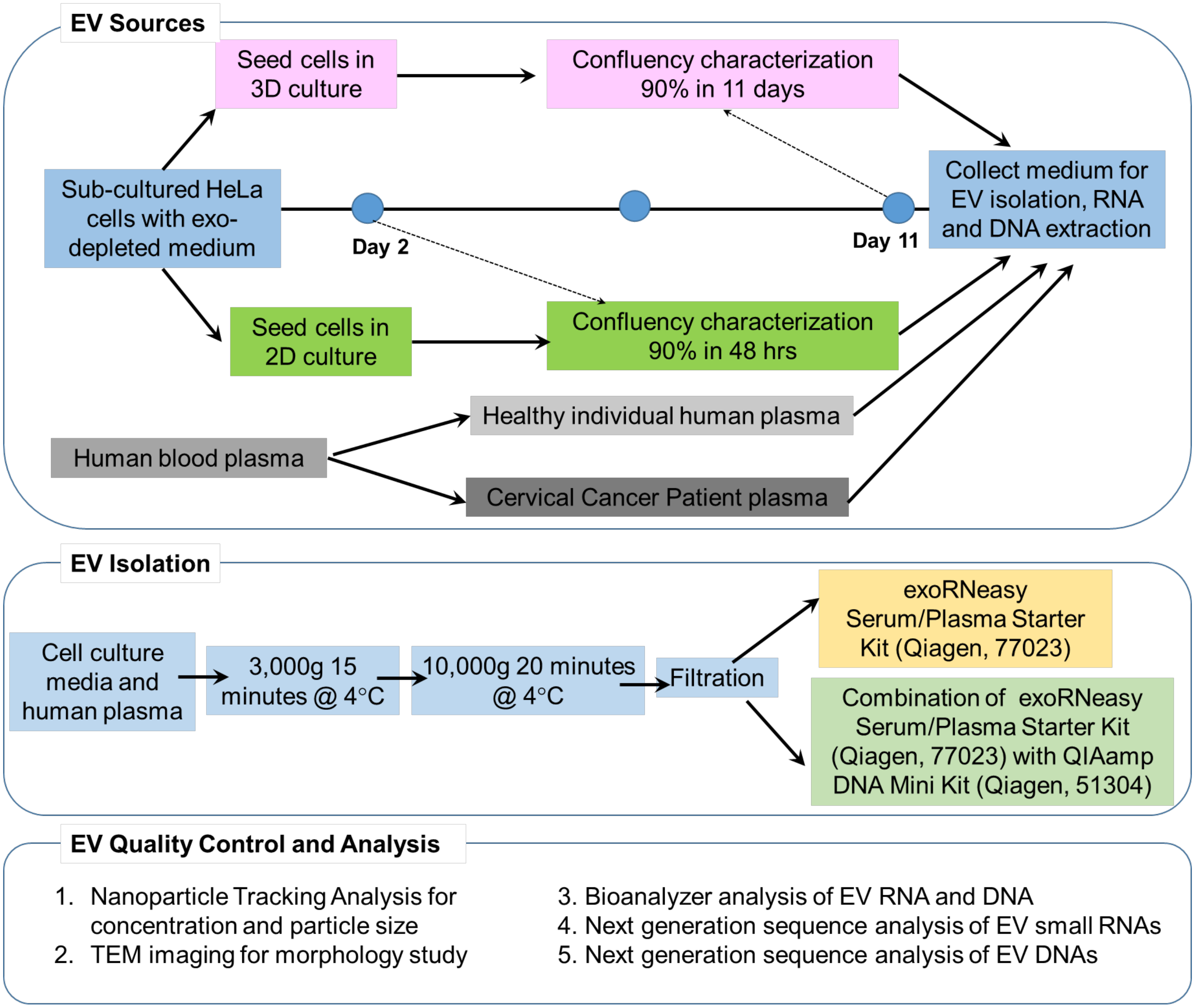


**Figure s1**. Scheme of the experimental flow for EV production, isolation, characterization, and data analysis. Quality control techniques (NTA, TEM, and Bioanalyzer) were performed on the isolated EVs prior to next-generation sequencing of small RNAs and DNAs.

**Characterization of 3D HeLa Cell Confluency Behavior:**

For characterizing 3D cell confluency behavior, we count the 3D cultured live cells at certain duration (6 hours after seeding, 2 days, 3 days, 5 days, 6 days and 8 days). The hydrogel can be broken down by pipetting up and down of culture for several times. Then the spheroids were suspended and transferred to 15 mL tube for rinsing with 2 mL of media twice. Mix 8 mL more media into the tube. Centrifuge at 250 g for 5 minutes. Use micro-pipette at 50 µL to remove all media from cell pellet but not to disturb the cell pellet. Resuspend the cell pellet in 50 µL Trypsin and mix well for 5 minutes, and then add 50 uL of Trypsin inhibitor in the mixture. Re-measure the total volume if possible (usually the volume will be 110 uL from residual media and cells). Add an equal amount of Trypan blue and use 10 uL to count cells (Cell counting chamber, Fisher).

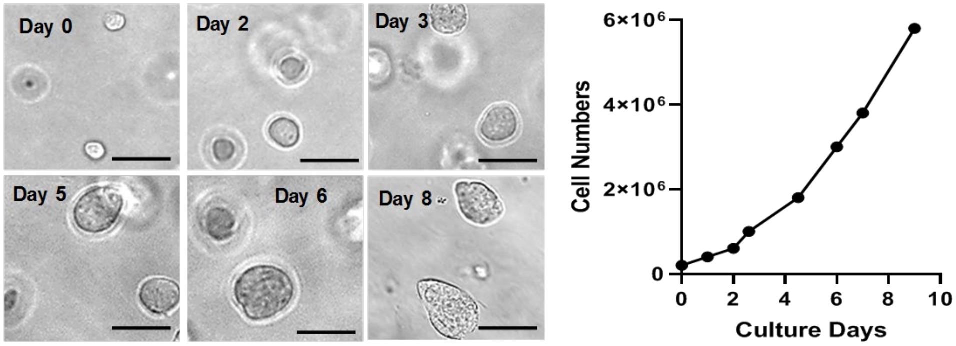


**Figure s2.** Time-dependent HeLa cell 3D culture. Cell culture conditions according to the method were described in the experimental section. Bright-field microscopic images (left) represent the 3D cell morphology cultured from 0 day (6 hours after seeding), 2 days, 3 days, 5 days, 6 days, to 8 days, respectively. The initial seeding density is ~$8\times10^4$ cells/mL. The scale bar is 80 µm. 3D cultured live HeLa cells are counted using the method described above along different culture durations for calibrating and estimating total cell numbers and growth rate (calibration curve in right hand).

**Quantitative Real-time PCR for Validating NGS Data**

TaqMan Advanced miRNA assays are delivered in a single tube containing the specific pre-formulated TaqMan Assay (TaqMan MGB probe, and forward and reverse primers). miR-Amp reagents are included in the TaqMan Advanced miRNA cDNA Synthesis Kit, and universal primers and master mix was used to uniformly increase the amount of cDNA for each target. We used six assay kits for selected miRNAs, including Hsa-miR-125b-1-3p, Hsa-miR-208a-5p, Hsa-miR-450b-3p, Hsa-miR-1229-3p, Hsa-miR-1284, Hsa-miR-1909-3p. We set up and run the real-time PCR instrument with the appropriate PCR thermal cycling conditions and selected the fast cycling mode for all instruments with 40 cycles.

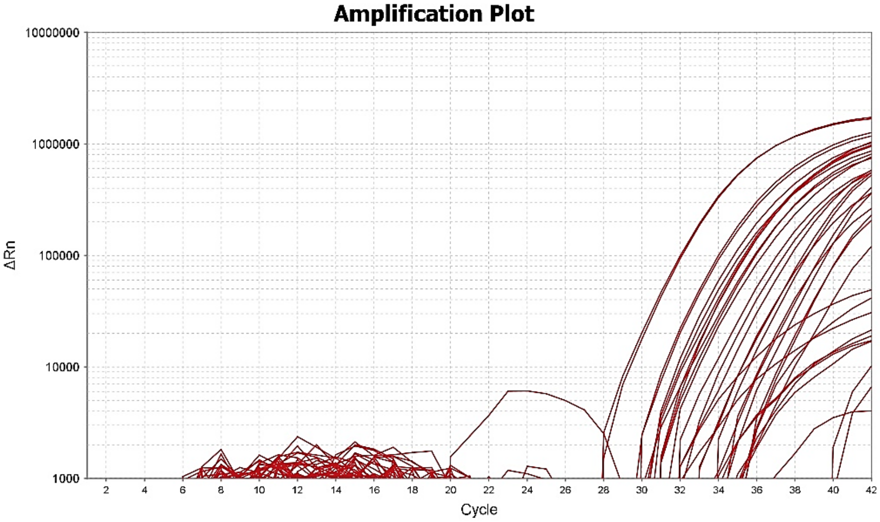


**Figure s3.** The amplification plots depict the qPCR cycles of six selected miRNAs with repeats.

**Table s1. Top pathways with 10-fold change for miRNAs significantly enriched in EVs from their parent cells (2D system)**

| *p*-value | Pathways | External IDs | Gene IDs |
|---|---|---|---|
| 4.05583E-16 | ABC transporters - Homo sapiens (human) | path:hsa02010 | 19; 22; 5825; 9429; 10349; 10350; 64241; 9619 |
| 7.94085E-13 | ABC-family proteins mediated transport | R-HSA-382556 | 5825; 23; 10350; 9619; 10349; 64241; 22 |
| 2.46769E-11 | ABC transporters in lipid homeostasis | R-HSA-1369062 | 10350; 5825; 10349; 9619; 64241 |

| | | | |
|---|---|---|---|
| 7.83492E-08 | Transport of small molecules | R-HSA-382551 | 10350; 64241; 19; 5825; 23; 22; 10349; 9619; 9429 |
| 1.46137E-05 | Nuclear Receptors in Lipid Metabolism and Toxicity | WP299 | 9619; 5825; 19 |
| 0.000543634 | Alanine Aspartate Asparagine metabolism | None | 16; 18 |
| 0.000838456 | Statin Pathway | WP430 | 64241; 19 |
| 0.001466377 | Fat digestion and absorption - Homo sapiens (human) | path:hsa04975 | 19; 64241 |
| 0.00217437 | Cholesterol metabolism - Homo sapiens (human) | path:hsa04979 | 19; 64241 |
| 0.002816538 | C21-steroid hormone biosynthesis and metabolism | C21-steroid hormone biosynthesis and metabolism | 9619; 64241 |
| 0.004099225 | Plasma lipoprotein assembly, remodeling, and clearance | R-HSA-174824 | 9619; 19 |
| 0.004334931 | Bile secretion - Homo sapiens (human) | path:hsa04976 | 9429; 64241 |

**Table s2. Top pathways with 10-fold change for miRNAs significantly enriched in EVs from their parent cells (3D system)**

| ***p*-value** | **Pathways** | **External IDs** | **Gene IDs** |
|---|---|---|---|
| 1.34641E-11 | ABC transporters - Homo sapiens (human) | path:hsa02010 | 64137; 4363; 19; 9429; 10350; 9619 |
| 9.24674E-09 | ABC-family proteins mediated transport | R-HSA-382556 | 23; 4363; 10350; 9619; 64137 |
| 1.5424E-06 | ABC transporters in lipid homeostasis | R-HSA-1369062 | 10350; 9619; 64137 |
| 1.16293E-05 | Transport of small molecules | R-HSA-382551 | 10350; 64137; 19; 23; 4363; 9619; 9429 |
| 3.43433E-05 | Methotrexate Pathway (Brain Cell), Pharmacokinetics | PA165816270 | 4363; 9429 |
| 3.43433E-05 | Paclitaxel Action Pathway | SMP00434 | 9429; 4363 |
| 3.43433E-05 | Docetaxel Action Pathway | SMP00435 | 9429; 4363 |
| 0.000126967 | Irinotecan Pathway | WP229 | 4363; 9429 |
| 0.000126967 | Taxane Pathway, Pharmacokinetics | PA154426155 | 4363; 9429 |
| 0.000126967 | Methotrexate Pathway, Pharmacokinetics | PA165816349 | 4363; 9429 |
| 0.000126967 | Irinotecan Pathway, Pharmacokinetics | PA2001 | 4363; 9429 |
| 0.000220692 | Doxorubicin Metabolism Pathway | SMP00650 | 4363; 9429 |
| 0.000248086 | Doxorubicin Pathway (Cancer Cell), Pharmacodynamics | PA165292163 | 4363; 9429 |
| 0.000248086 | Lamivudine Metabolism Pathway | SMP00649 | 4363; 9429 |
| 0.000248086 | Lamivudine Pathway, Pharmacokinetics/Pharmacodynamics | PA165860384 | 9429; 4363 |
| 0.000373399 | Doxorubicin Pathway, Pharmacokinetics | PA165292177 | 4363; 9429 |
| 0.000408644 | Irinotecan Action Pathway | SMP00433 | 4363; 9429 |
| 0.000408644 | Irinotecan Metabolism Pathway | SMP00600 | 4363; 9429 |
| 0.000445447 | Pathway_PA165986194 -need delete | PA165986194 | 4363; 9429 |
| 0.000445447 | Acetaminophen Pathway, Pharmacokinetics | PA165986279 | 4363; 9429 |
| 0.0006988 | Acetaminophen Metabolism Pathway | SMP00640 | 4363; 9429 |

| | | | |
|---|---|---|---|
| 0.000952126 | Nuclear Receptors in Lipid Metabolism and Toxicity | WP299 | 9619; 19 |
| 0.003656373 | Plasma lipoprotein assembly, remodeling, and clearance | R-HSA-174824 | 9619; 19 |

**Reference:**


1. Gheinani AH, Vogeli M, Baumgartner U, Vassella E, Draeger A, Burkhard FC, Monastyrskaya K: **Improved isolation strategies to increase the yield and purity of human urinary exosomes for biomarker discovery**. *Sci Rep* 2018, **8**(1):3945.
